# Modelling the effect of matrix metalloproteinases in dermal wound healing

**DOI:** 10.1101/2023.05.24.542101

**Authors:** Sonia Dari, Nabil T. Fadai, Reuben D. O’Dea

## Abstract

With over 2 million people in the UK suffering from chronic wounds, understanding the biochemistry and pharmacology that underpins these wounds and wound healing is of high importance. Chronic wounds are characterised by high levels of matrix metalloproteinases (MMPs), which are necessary for the modification of healthy tissue in the healing process. Overexposure of MMPs, however, adversely affects healing of the wound by causing further destruction of the surrounding extracellular matrix. In this work, we propose a mathematical model that focuses on the interaction of MMPs with dermal cells using a system of partial differential equations. Using biologically realistic parameter values, this model gives rise to travelling waves corresponding to a front of healthy cells invading a wound. From the arising travelling wave analysis, we observe that deregulated apoptosis results in the emergence of chronic wounds, characterised by elevated MMP concentrations. We also observe hysteresis effects when both the apoptotic rate and MMP production rate are varied, providing further insight into the management (and potential reversal) of chronic wounds.

## 1 Introduction

Wound healing is a physiological response to injury of tissue involving the coordinated interactions of many cell types and biochemical agents [1, 2]. In individuals such as diabetes patients, so-called ‘chronic wounds’ may persist, which require medical treatments to promote healing [3, 4]. There are approximately 2.2 million people in the UK suffering from chronic wounds, costing the NHS over £5 billion per year [5]. These wounds last, on average, 12–13 months and recur in 60%–70% of individuals, potentially leading to the loss of function [6]. Improved understanding of the biochemical mechanisms underpinning chronic wounds is therefore crucial to support the development of new treatments.

The healing of wounds involves the complex interplay of various cell types and their mediators and cytokines [7]. A wound is defined as damage to the skin which is comprised of two layers: the epidermis and the dermis. The epidermis is the outermost layer and is responsible for protecting against infection, while the dermis is the innermost layer and provides tensile strength for the skin by means of the dermal extracellular matrix (ECM) [7, 8]. The process of wound healing can be categorised into four successive stages: haemostasis, inflammation, proliferation and remodelling [1]. A wound that is able to complete these four stages in a well-coordinated manner is defined as an acute wound; conversely, a wound that spends a prolonged time in any of the four stages is defined as a chronic wound [3]. The inflammation and proliferation stages of wound healing involve the production of matrix metalloproteinases (MMPs) by fibroblasts [9]. MMPs are responsible for lysing protein components of the ECM, allowing for fibroblast migration within the ECM and, as a result, the proliferation of cells and modification of tissue [3]. Experimental data has shown that there is increased expression of MMPs at the edge of a wound in all stages of the healing of an acute wound [10]. It is essential to have a well regulated concentration of MMPs during this process: if MMP levels are too low, this leads to uncontrolled ECM production and can cause issues such as hypertrophic scarring or dermal fibrosis [11]. If MMP levels are too high, chronic wounds are observed, as the overexposure of MMPs results in the increased degradation of the ECM [12]. One such cause for these elevated MMP levels is deregulated apoptosis of dermal tissue, which may be caused by uncontrolled blood sugar levels in diabetes patients [13, 14].

Mathematical modelling of the wound healing process can provide a framework to understand the behaviour of chronic wounds, potentially to direct treatments and improve the quality of life of patients. Many mathematical models have been developed to describe the wound healing process, each focusing on specific mechanisms of interest. For example, agent–based models (ABMs) can emulate the individual-level stochastic nature of the biological processes involved in wound healing [15–20]. Although ABMs are able to detail specific properties of individual cells, using them to model dynamics of wound healing on a tissue level may be computationally infeasible. We therefore will consider continuous models of wound healing, which typically use partial differential equations (PDEs) and present opportunities for a range of analytical techniques for further study [7].

Continuous mathematical models of both dermal and epidermal wound healing typically take the form of reaction-diffusion systems, extending the work of Sherratt and Murray [21] that examined the interaction of epidermal cells with an representative chemical acting as a regulator of mitosis. Their model has subsequently been developed to consider additional chemical agents involved in epidermal wound healing, such as the role of epidermal growth factor in corneal wound healing and the role of keratinocyte growth factor in epidermal wound healing [22–25]. Contrastingly, other models of dermal wound healing consider the role of fibroblasts in restoring the ECM in response to various mechanisms at different stages in the wound healing process. Many of these models consider the roles of angiogenesis and consider the role of oxygen transport in wound healing [7, 26–30], while others consider the process of ECM restoration by fibroblasts in response to growth factors [31, 32], the crosstalk of the epidermis and the dermis to simulate the simultaneous healing of both layers [33, 34] and the incorporation of collagen fibre orientation due to cell movement [35–39]. Travelling wave solutions are often observed in these mathematical models, agreeing with the experimental evidence of wound healing assays [40, 41]. However, these models are often limited to the healing of acute wounds.

In this work, we consider the role that MMPs play in wound healing, with particular consideration given to chronic wounds. We propose a two-component reaction-diffusion model that describes the interaction of MMPs with dermal tissue, in which the restoration of the ECM is affected by MMP concentration levels. Using biologically realistic parameter values, this model gives rise to travelling waves corresponding to a front of healthy cells invading a wound. From the arising travelling wave analysis, we observe that deregulated apoptosis results in the emergence of chronic wounds, characterised by elevated MMP concentrations. We also observe hysteresis effects when both the apoptotic rate and MMP production rate are varied, providing further insight into the management (and potential reversal) of chronic wounds.

We develop our mathematical model in Section 2. Direct numerical simulations indicate the existence of travelling wave solutions, motivating a travelling wave analysis in Section 3. We then consider the effect of deregulated apoptosis on the healing of a wound via bifurcation and phase plane analysis in Section 4. Finally, we consider the effects of elevated MMP production levels on the healing of a wound in Section 6, before discussing our results in Section 7.

## 2 Model Development

We begin by presenting a mathematical formulation to describe MMP concentration dynamics in the presence of a wound. Elevated levels of MMPs play a key role in the persistence of chronic wounds; in light of this, we focus on constructing a reaction-diffusion model of wound healing that incorporates the interaction between MMPs with dermal cells during the wound healing process. For simplicity, we consider a wound with a one-dimensional Cartesian geometry, where *x* represents the longitudinal distance across the wound, which evolves over time *t*. This model consists of two populations within a wound: the dermal cell density, denoted as *n*(*x, t*), and the MMP concentration, denoted as *m*(*x, t*). We take *n*(*x, t*) to incorporate the actions of both fibroblasts and ECM for the sake of simplicity, since fibroblasts are responsible for creating the components of the ECM.

We assume that dermal cells undergo mitosis with intrinsic growth rate σ and carrying capacity *K*_*n*_, noting that *n* = 0 corresponds to the complete absence of cell tissue. As mentioned previously, MMPs can assist in the healing of the wound up to a threshold concentration, denoted here as *m*_thresh_, above which they adversely affect healing of the wound by degrading the surrounding ECM. This effect is incorporated in the mitosis process by means of a function *f*(*m*). Furthermore, dermal cells undergo apoptosis with rate *δ*_*n*_ and dermal cell motion within the wound is represented by linear diffusion with effective diffusivity *D*_*n*_.

MMP production by fibroblasts, denoted by a recruitment function *g*(*n*), occurs in response to inflammatory chemicals which are found in abundance in a wound, and thus is assumed here to be proportional to the absence of healthy tissue. MMPs undergo natural decay with rate *δ*_*e*_ and diffuse through the wound with diffusivity *D*_*m*_. These aforementioned processes are shown in Figure 1. Combining these modelling elements provides the following system of PDEs:

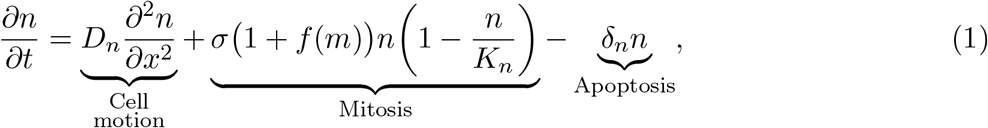

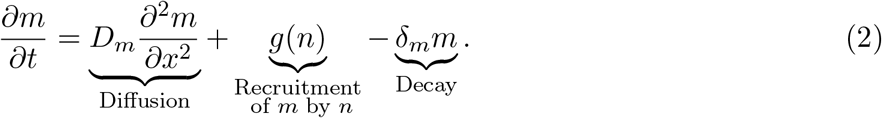

We now discuss the functional forms of *f*(*m*) and *g*(*n*) appearing in the system (1)–(2). The function *f*(*m*) in (1) represents the contribution to wound healing by MMPs and is defined to be an increasing function of *m* until *m* = *m*_thresh_, at which *f*(*m*) = *f*_max_, the maximum factor by which mitosis can be enhanced due to the presence of MMPs. Larger concentrations of MMPs hinder the wound healing process and thus, for *m* larger than *m*_thresh_, *f*(*m*) is a decreasing function of *m*. A suitable choice of *f*(*m*)is therefore

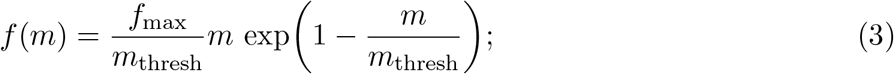

a schematic of *f*(*m*) is shown in Figure 2(a). We note that *f*(*m*) and hence the parameter *f*_max_ are dimensionless. For elevated levels of MMPs, *f*(*m*) decreases to zero and the mitosis term reduces to logistic growth with zero healing contribution due to MMPs. One could also allow *f*(*m*) to take negative values for large *m*, representing a negative healing contribution to the ECM. We have considered a functional form of this type in Appendix A, where we show that such a choice gives rise to largely similar qualitative features to those produced by the function form of *f*(*m*) given in (3). Given these similarities, we restrict attention to the choice of *f*(*m*) as in (3) as this choice results in simpler analytic computations.

**Figure 1:**
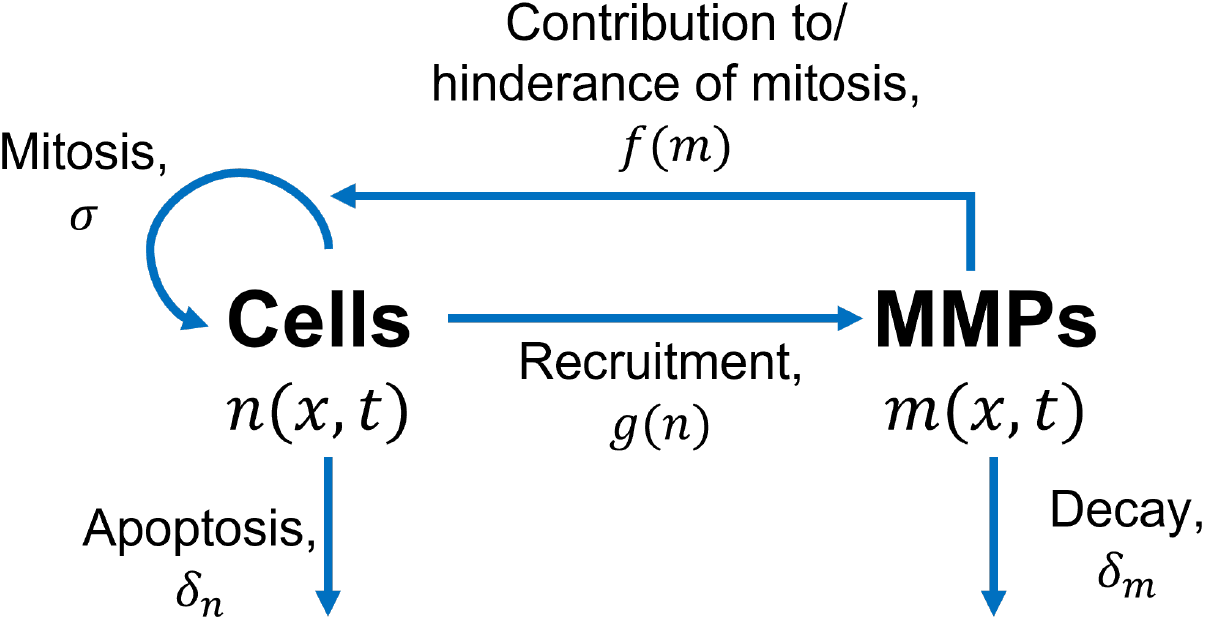
Schematic of the interactions of dermal cells with MMPs. Labelled rates are defined as in the system (1)–(2).

**Figure 2:**
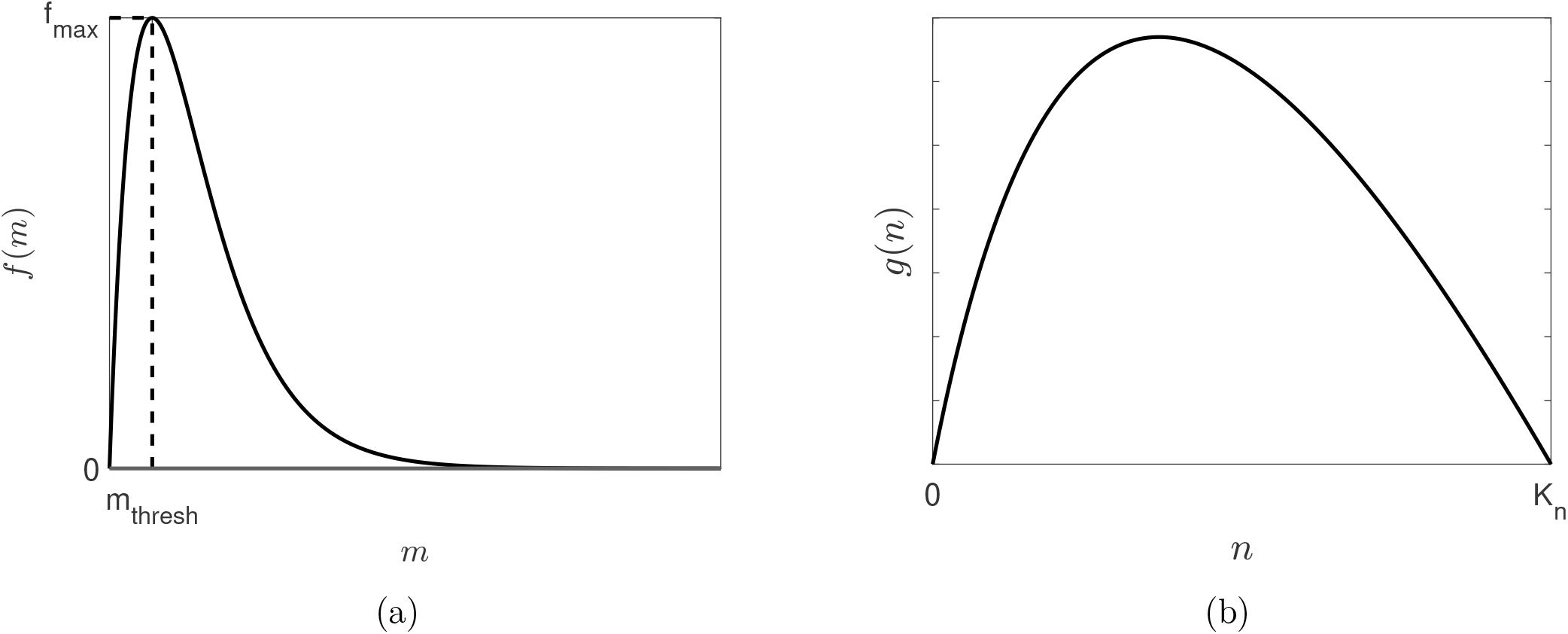
Schematic functional forms of (a) the healing contribution by MMPs function, *f*(*m*) in (3), and (b) the MMP production function, *g*(*n*) in (4).

The recruitment function *g*(*n*) in (2) represents the production of MMPs by the cells in pro-portion to the absence of healthy tissue. Consequently, we can represent *g*(*n*) as

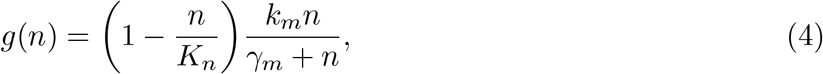

where *k*_*m*_ is the maximum reaction rate and *γ*_*m*_ is a Michaelis–Menten constant [42]. A schematic of *g*(*n*) is shown in Figure 2(b).

### 2.1 Initial Conditions and Boundary Conditions

The initial conditions of the system (1)–(2) are chosen as follows to emulate a wound starting at *x* = λ, with a small concentration of MMPs, *η*, at the wound edge:

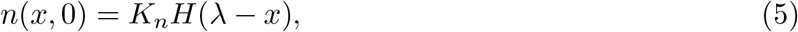

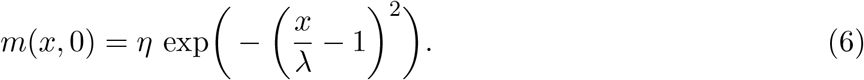

In (5), *H* is the Heaviside function while *n* = *K*_*n*_ corresponds to the cells being at carrying capacity. Assuming the wound to be symmetric about its centre and that the tissue is fully healed far away from the wound, we adopt zero-flux boundary conditions on both endpoints of the finite domain of length *L*, i.e:

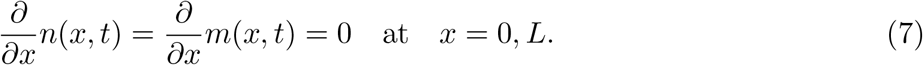

### 2.2 Nondimensionalisation

In this section, we non-dimensionalise the system (1)–(7). We are interested in the dynamics of the system on the timescale of mitotic rate of dermal cells and therefore introduce the following dimensionless variables:

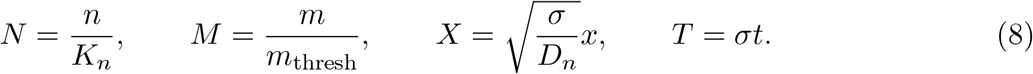

With these scalings, we obtain the following dimensionless equations:

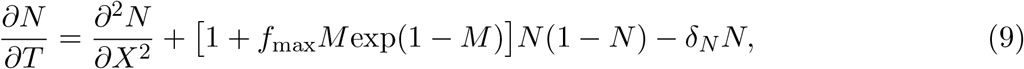

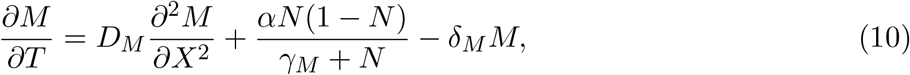

where

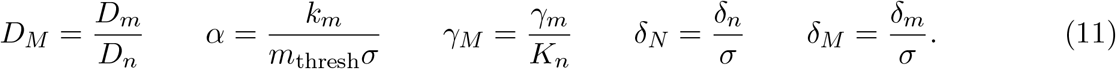

The dimensionless initial and boundary conditions are:

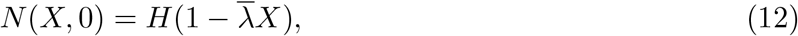

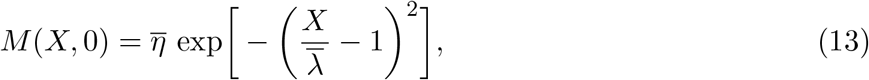

and

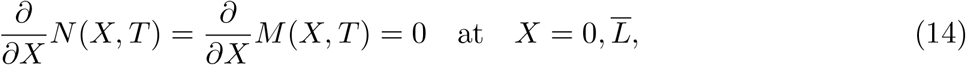

where 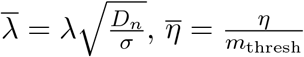 and 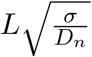.

We define the dimensionless system (9)–(14) as the ‘wound healing model’ and we note that 0 ≤ *N*(*X, T*) ≤ 1, since we do not expect the cell density to exceed its carrying capacity.

Suitable values for the dimensional parameters appearing in (1)–(4) are given in Table 1; we remark that these parameters correspond to ‘normal’, or healthy, wound healing and, unless otherwise stated, are employed in all simulations. These parameter values are used to calculate the dimensionless parameter values in (11)–(14) and are given in Table 2.

**Table 1:** Values for the dimensional parameters appearing in (1)–(4).

| Parameter | Description | Value | Reference |
| --- | --- | --- | --- |
| $D_n$ | Diffusion coefficient of cells | $6.12 \times 10^{-5} \text{ mm}^2 \text{ h}^{-1}$ | [30] |
| $D_m$ | Diffusion coefficient of MMPs | $9 \times 10^{-4} \text{ mm}^2 \text{ h}^{-1}$ | [43] |
| $\sigma$ | Mitotic rate of cells | $0.0385 \text{ h}^{-1}$ | [44] |
| $K_n$ | Carrying capacity of cells | $1250 \text{ cells mm}^{-2}$ | [44] |
| $k_m$ | Maximal rate of MMP production by cells | $4 \text{ pg}/\mu\text{g h}^{-1}$ | Estimate |
| $\gamma_m$ | Density of cells at which half of $k_e$ is attained | $625 \text{ cell mm}^{-2}$ | Estimate |
| $f_{\max}$ | Maximum factor with which mitosis can be enhanced due to the presence of MMPs | 50 | Estimate |
| $\delta_m$ | MMP decay rate | $0.009 \text{ h}^{-1}$ | [45] |
| $\delta_n$ | Cell apoptosis rate | $0.005 \text{ h}^{-1}$ | [44] |
| $m_{\text{thresh}}$ | Threshold concentration of MMPs at which maximum healing contribution $f_{\max}$ is attained | $7 \text{ pg}/\mu\text{g}$ | [46] |

**Table 2:** Values of the dimensionless parameter values appearing in (9)–(14) calculated using the values given in Table 1.

| Parameter | Value |
| --- | --- |
| $D_M$ | 14.706 |
| $\alpha$ | 14.842 |
| $f_{\max}$ | 50 |
| $\delta_N$ | 0.130 |
| $\delta_M$ | 0.234 |
| $\gamma_M$ | 0.5 |
| $\bar{\eta}$ | 0.1 |
| $\bar{\lambda}$ | 3 |
| $\bar{L}$ | 200 |

### 2.3 Model Simulations

In order to obtain numerical solutions of the wound healing model, we use the method of lines and discretise on a one-dimensional spatial domain, using second order finite differences to approximate spatial derivatives. We then integrate the resulting system of time–dependent ODEs using MATLAB’s ODE15s solver.

An example numerical simulation of the wound healing model is shown in Figure 3. As seen in Figure 3(a), the cell density *N*(*X, T*) evolves into a travelling wave where a front of tissue of density *N* ≈ 1 is ‘invading’ the wound (*N* = 0). This state of *N* ≈ 1 is close enough to unity to be considered as a healed state; we elaborate on the classification of a healed state further in Section 3. Since we observe a travelling wave where a healed front of tissue invades the wound, this scenario corresponds to a wound healing to completion. In Figure 3(b), we also observe the formation of a travelling wave for *M*(*X, T*) in which a ‘spike’ in MMP concentration is found at the edge of the healing wound. These qualitative phenomena are in agreement with biological literature, which suggests the increased expression of MMPs at the edge of an acute would at all stages in the healing process [10].

**Figure 3:**
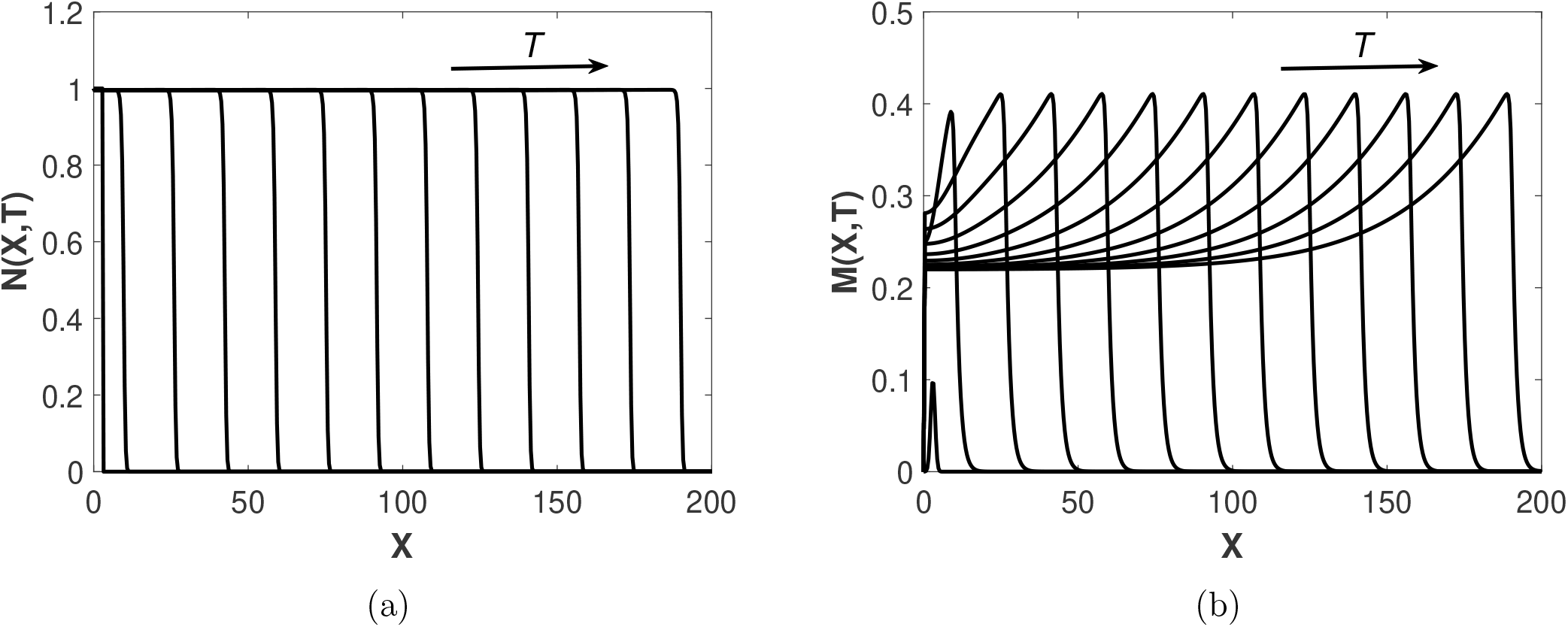
Simulation of the wound healing model (9)–(14) at regular time intervals *T* = 2 with parameter values as in Table 2. (a) Evolution of cell density *N*(*X, T*) across the spatial domain, (b) Evolution of MMP concentration *M*(*X, T*) across the spatial domain.

## 3 Steady State and Travelling Wave Analysis

From the previous section, we observe that the wound healing model gives rise to travelling wave solutions. Therefore, in this section we conduct a travelling wave analysis of the model. To determine the far-field states of the travelling waves, we first examine the uniform steady-states of (9)–(10). In particular, the invading far-field state for *N*(*X, T*) will indicate whether or not a wound is healing to completion. The uniform steady states 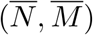 of (9)–(10) satisfy the following:

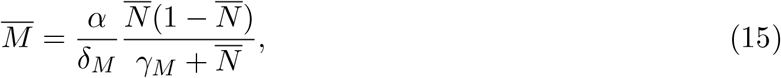

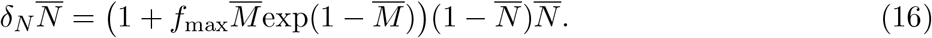

Combining (15)–(16), we find that all non-trivial steady states may be written in terms of the single variable 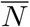 :

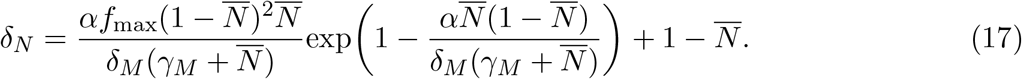

Using the fimplicit function in MATLAB, we determine a numerical solution for 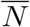 and subsequently 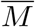 using equation (15).

It is clear from (16) that we always have the trivial steady state, i.e. 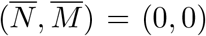 for all positive parameters. As discussed in Section 2.3, we characterise a wound as healing to completion when a far-field state of *N* ≈ 1 is achieved. This is because for *δ*_*N*_ = 0, 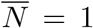 satisfies (16). For non-zero *δ*_*N*_, however, 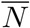 does not attain the value one, since apoptosis of cells is continually occurring. For small *δ*_*N*_, i.e. values corresponding to healthy biological functioning, (16) provides 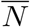 close to unity, and we hence consider this state to describes healed tissue. Additionally, if an unhealed or partially healed steady-state of 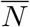, i.e. not close to one, invades a state corresponding to damaged tissue, we characterise this as a chronic wound.

The stability of the uniform steady-states 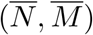 is given by linear stability analysis of the analogous spatially-independent form of the wound healing model:

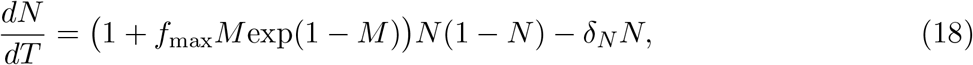

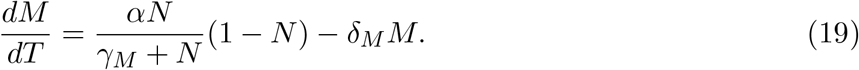

This stability analysis (detailed in Appendix B) is used to determine the stability of the branches of bifurcation diagrams which we examine in Sections 4 and 6.

In order to verify the existence of travelling solutions to the wound healing model, we consider a travelling wave analysis. Employing the travelling wave coordinates *ξ* = *X* − *cT* ∈ ℝ, where *c* is the wave speed, the wound healing model reduces to the boundary value problem (BVP):

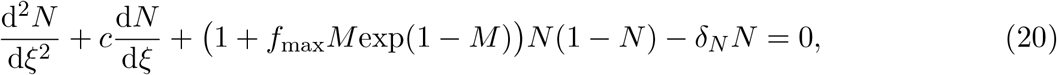

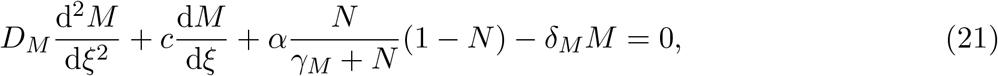

with boundary conditions

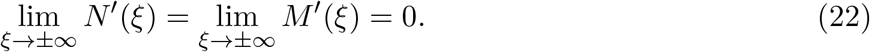

We note that when considering solutions in travelling wave coordinates, the stability of 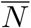 and 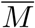 in the bifurcation diagrams in Figure 5 and 9 are swapped, i.e. stable branches are unstable in travelling wave coordinates and vice versa, due to the change of variables *ξ* = *X* − *cT*.

Using MATLAB’s BVP5c solver, we present numerical simulations of the BVP (20)–(22) in Figure 4 with parameter values given in Table 2, which verifies the simulations of the wound healing model in Figure 3. We are also able to use BVP5c to obtain a nondimensional wavespeed of *c* = 9.6, which translates to a dimensional wavespeed of *c*_dim_ = 1.504 × 10^−2^mm h^−1^. This wavespeed is of the correct order of magnitude observed from wound healing assays from [40].

**Figure 4:**
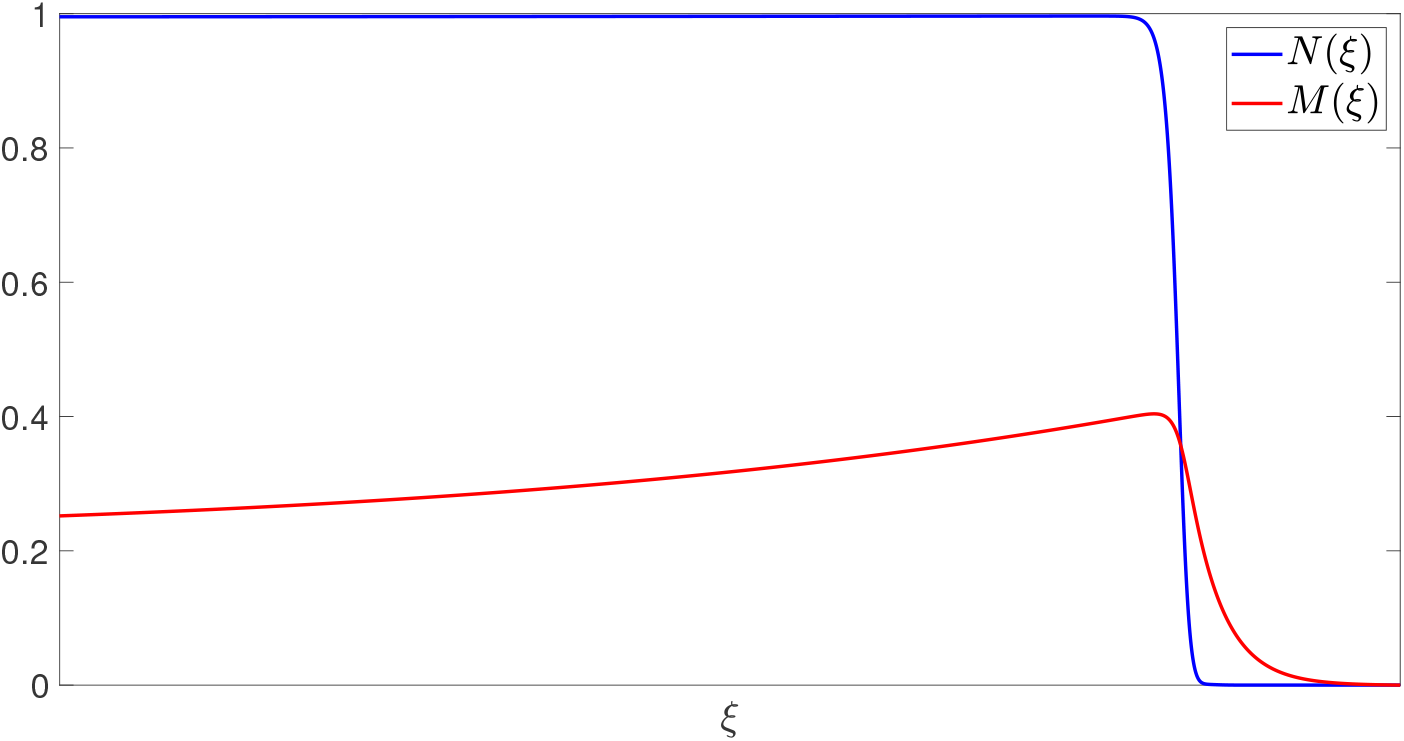
Numerical solution of the boundary value problem (20)–(22) obtained using MATLAB’s BVP5c solver with parameter values as in Table 2.

**Figure 5:**
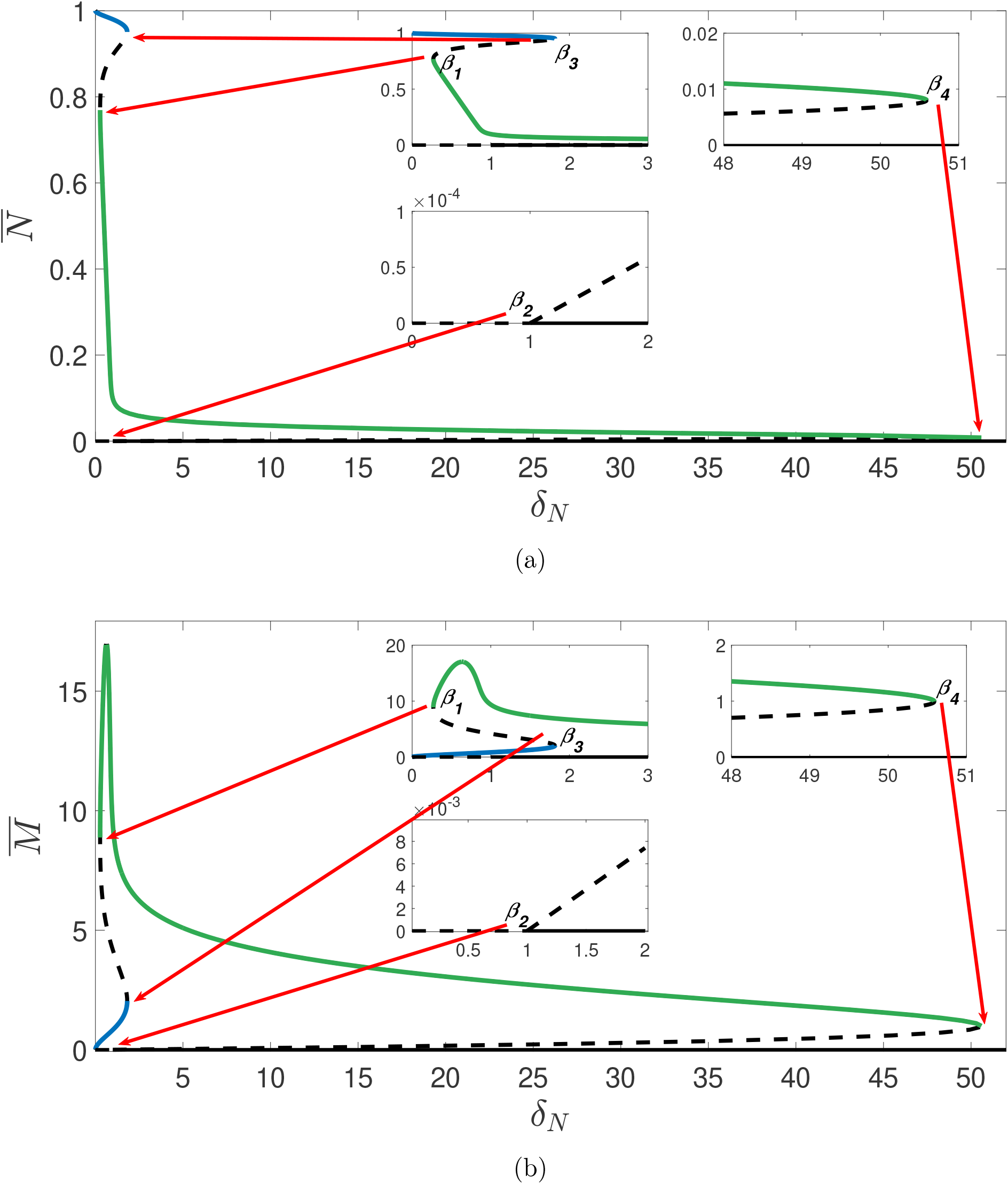
Bifurcation diagrams of (15)–(17) showing the steady-states (a) 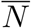 and (b) 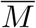 under variation of *δ*_*N*_. Other parameter values are given in Table 2. The bifurcation diagrams show four bifurcation points *β*_1_, *β*_2_, *β*_3_ and *β*_4_ separating five regimes *I*_1_, *I*_2_, *I*_3_, *I*_4_ and *I*_5_. Dashed lines represent unstable branches and solid lines represent stable branches. The stable branches are colour-coded such that the green branch of steady states of 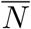 in (a) correspond to the green branch of steady states of 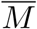 in (b), and similarly for blue branches.

## 4 Deregulated Apoptosis

As mentioned previously, a potential cause for the emergence of chronic wounds is deregulated apoptosis. In this section, we examine the behaviour of the wound healing model under variation of the apoptotic rate of cells, *δ*_*N*_. Figure 5 shows bifurcation diagrams for *N*(*X, T*) and *M*(*X, T*) against *δ*_*N*_; the stability of the branches is determined by linear stability analysis that is detailed in Appendix B.

Inspection of the bifurcation diagrams in Figure 5 show that we have four bifurcation points that separate five parameter regimes, in which solution behaviours of the wound healing model differ qualitatively. The intervals for each regime are denoted as follows:

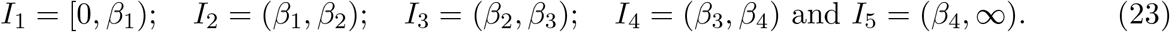

The positions of the interval boundaries *β*_1_ *< β*_2_ *< β*_3_ *< β*_4_ and the method used to calculate them are given in Appendix C. While each interval is discussed in greater detail in subsequent subsections, we begin with a brief overview of the qualitative behaviours observed in each region.

For *δ*_*N*_ ∈ *I*_1_, we have two steady-states for 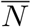 : a state approaching unity and the trivial state. As discussed in Section 2.3, this parameter regime corresponds to a wound healing to completion, with an increased expression of MMPs at the wound edge. For *δ*_*N*_ ∈ *I*_2_, *I*_3_, it is unclear from the bifurcation diagram whether or not a wound will heal to completion, due to the existence of other non-trivial steady states. The bifurcation diagram for 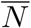 indicates that we have 3 and 4 nontrivial states for *I*_2_ and *I*_3_ respectively: with one being a healed state 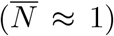, and the others being partially healed states. At *δ*_*N*_ = *β*_3_, we observe a saddle node bifurcation point at which the healed state is destroyed. For *δ*_*N*_ ∈ *I*_4_, there are two non-trivial states for 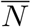, both being unhealed states which indicates that for *δ*_*N*_ ∈ *I*_4_, a wound will not heal to completion and hence a chronic wound will persist. Finally, for *δ*_*N*_ ∈ *I*_5_ only the trivial state exists.

### 4.1 *δ*_*N*_ ∈ *I*_1_

The bifurcation diagram in Figure 5 indicates that we have two steady-state solutions, with one being the trivial state. The other state is 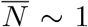 at leading order, which can be verified using a regular perturbation analysis by considering the case where *δ*_*N*_, *δ*_*E*_ ≪ 1. As discussed in Section 3, this regime describes the invasion of a wound by a healed front of cells and thus the healing of an acute wound. These features are reflected in direct simulation of the full system in Figure 6(a), for which *δ*_*N*_ ∈ *I*_1_. We also note that travelling waves for *M*(*X, T*) in Figure 6(b) display an increased expression of MMP concentration at the wound edge, which is in agreement with the biological literature as discussed in Section 1. From the phase plane diagram in Figure 6(c), we see that regardless of initial conditions, trajectories in (*N, M*) space always arrive at the healed state, 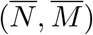. Therefore, for *δ*_*N*_ ∈ *I*_1_ and other parameter values as in Table 2, a wound will always heal to completion.

**Figure 6:**
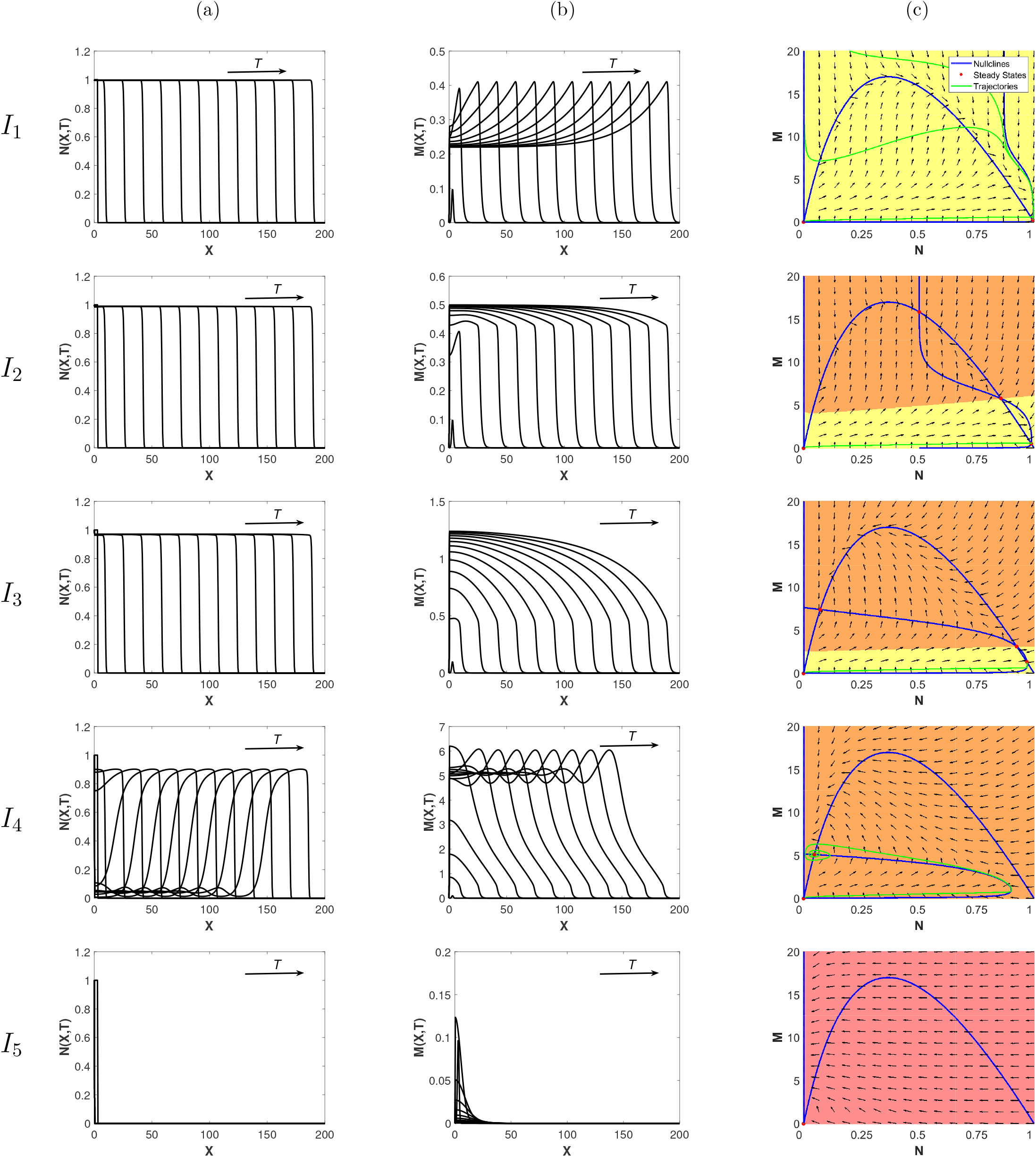
Direct simulations of the wound healing model, for various *δ*_*N*_ values for (a) *N*(*X, T*) and (b) *M*(*X, T*). The phase plane diagrams in (c) correspond to the spatially independent ODE system (18)–(19). Blue lines represent nullclines, green lines indicate representative trajectories of the ODE system and red dots represent the steady states. The yellow highlighted section of the domain represents the set of all the initial conditions (ICs) in the (*N, M*) phase space whose trajectories go to the healed state, the orange section represents the ICs whose trajectories go to an unhealed state and the red section represents the ICs whose trajectories go to the zero state. All other parameter values are as given in Table 2. We note that for *I*_2_ and *I*_3_, only the fully healed travelling waves (obtained using initial conditions as in (12)–(13)) are shown in (a) and (b). We also note that the *M* -axis changes in each row.

### 4.2 *δ*_*N*_ ∈ *I*_2_, *I*_3_

For *δ*_*N*_ ∈ *I*_2_, *I*_3_, the bifurcation diagram in Figure 5 shows that we have 3 and 4 nontrivial states, respectively. The phase plane diagrams in Figure 6(c) indicate that the initial state determines whether or not the wound heals to completion, with high initial concentrations of MMPs resulting in the emergence of a chronic wound. Contrastingly, if a small concentration of MMPs is initially present at the wound edge, a wound can heal to completion (Figures 6(a)). This suggests that baseline levels of MMP must be kept low in order to minimise risk of the persistence of a chronic wound. The simulations for *δ*_*N*_ ∈ *I*_2_, *I*_3_ exhibit behaviour similar to that for which *δ*_*N*_ ∈ *I*_1_, demonstrating the healing of an acute wound.

### 4.3 *δ*_*N*_ ∈ *I*_4_

For *δ*_*N*_ ∈ *I*_4_, all trajectories arrive at one of two unhealed states for all initial conditions (Figure 6(c)). These states have complex eigenvalues, characterised by spirals in the phase plane diagram and we therefore expect a travelling wave corresponding to a chronic wound with fluctuating MMP levels about its nonzero state. This is verified by the simulations in Figure 6(a) and 6(b), where we observe travelling waves with qualitatively different features to those observed in previously discussed intervals.

This change in the qualitative features of the travelling waves corresponds to a saddle-node bifurcation point at *δ*_*N*_ = *β*_3_ observed in the bifurcation diagrams in Figure 5. At this bifurcation point, the branch of healed states 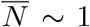, shown in blue (and the corresponding branch for 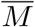) is destroyed. This loss of a healed state of 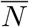 indicates that the invading state in the travelling wave simulations will transition to an unhealed state, which is verified by Figure 6(a) where we also observe increased density of dermal tissue at the edge of the wound. As discussed in Section 3, we characterise this as a chronic wound. The emergence of a chronic wound also corresponds to the invading state of 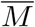 in the travelling wave simulations transitioning to a larger value as shown in Figure 6(b), suggesting that elevated MMP concentrations prevent the complete healing of the wound. These results are consistent with the biological literature in that deregulated apoptosis is a prominent cause of the emergence of chronic wounds, which are characterised by having raised MMP concentrations.

### 4.4 *δ*_*N*_ ∈ *I*_5_

For *δ*_*N*_ ∈ *I*_5_, we find that *N* and *M* both tend to zero for all initial conditions. This collapse of the travelling wave is expected in the case where apoptosis of dermal cells is very high, as cells die faster than they can reproduce.

## 5 Multistability and Hysteresis

As discussed in Section 1, a potential cause for the emergence of chronic wounds is deregulated apoptosis which may be caused by uncontrolled blood sugar levels in diabetes patients. In this section, we therefore examine how the apoptotic rate, *δ*_*N*_ controls the transition from a healthy to a chronic state. We also investigate whether or not a chronic wound may be reversed if the apoptotic rate were to be regulated, for example, by regulating blood sugar levels. As discussed in Section 4.3, a saddle-node bifurcation point occurs at *δ*_*N*_ = *β*_3_, which results in the emergence of a chronic wound, where high concentrations of MMPs prevent the wound healing to completion. The subcritical nature of this bifurcation has important implications for the solution behaviour, which we demonstrate in direct simulations of the wound healing model by allowing *δ*_*N*_ in (9) to slowly increase in time:

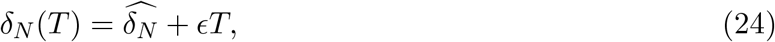

where 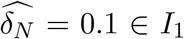. As we see in Figure 7, the travelling wave for *N*(*X, T*) transitions from a completely healed wound to a chronic wound with elevated MMP concentrations, which verifies the results outlined in Section 4.3.

**Figure 7:**
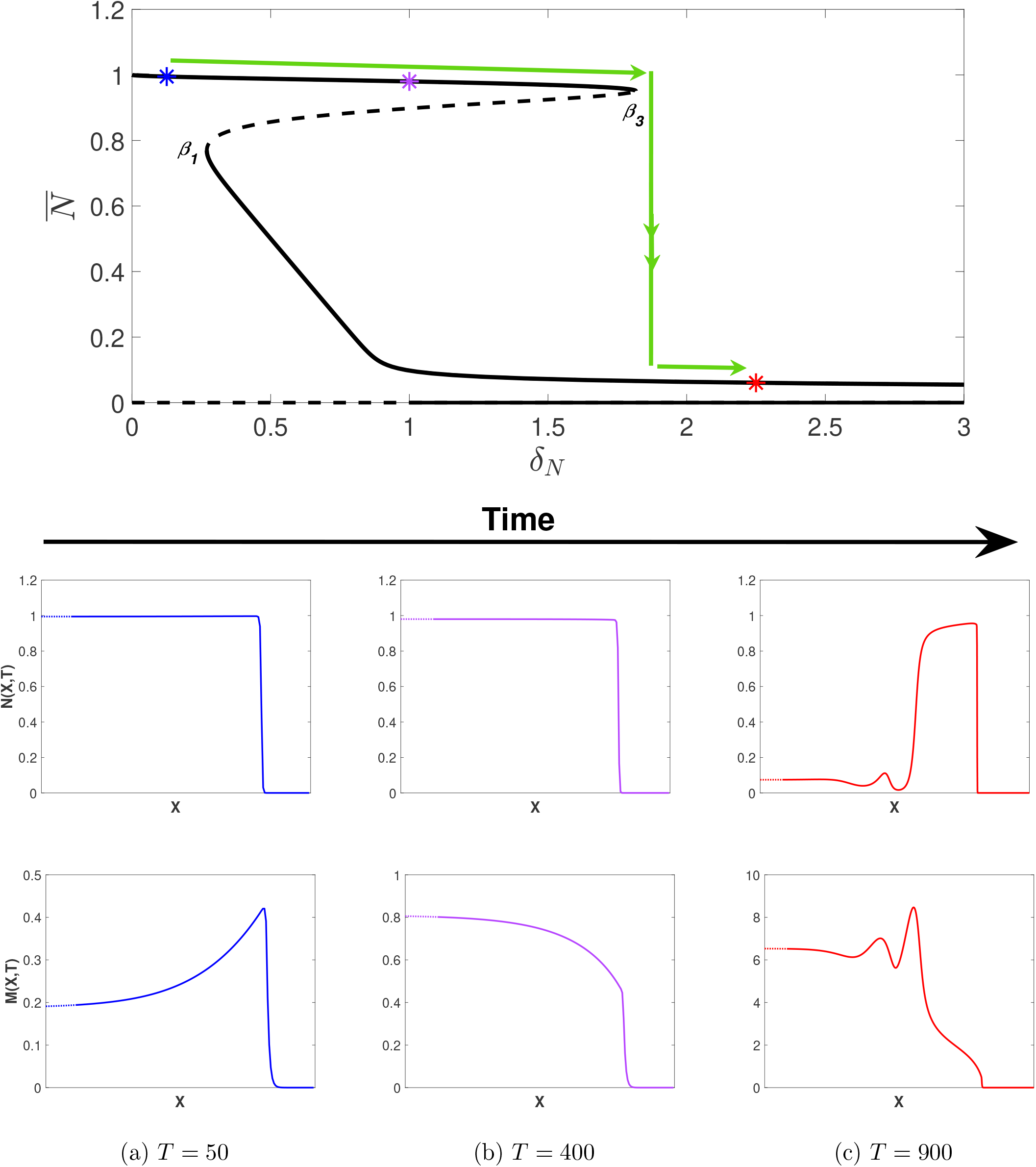
Simulations of the wound healing model with (24) applied to (9), i.e. *δ*_*N*_ increasing in time with 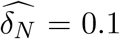 and *ϵ* = 0.01. Other parameter values are as given in Table 2. Note that the *X*-axis has been trimmed to highlight the qualitative features of each travelling wave (dotted lines). The bifurcation diagram is as given in Figure 5(a) for *δ*_*N*_ ∈ [0, 3] and coloured asterisks represent the *δ*_*N*_ (*T*) values at the time points given in (a), (b) and (c). Note that all travelling waves shown connect to the zero state.

As mentioned in Section 4.3, a saddle-node bifurcation point also occurs at *δ*_*N*_ = *β*_1_. As such, we consider the case of decreasing *δ*_*N*_ to investigate the implications for the solution behaviour upon decreasing *δ*_*N*_ below *β*_1_ from a regime in which a chronic wound persists. We take a value of 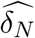 such that 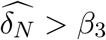 (to ensure the persistence of a chronic wound), and allow *δ*_*N*_ in (9) to slowly decrease in time via (24) by changing *ϵ* with −*ϵ*. In Figure 8, we take 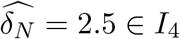 and simulate the wound healing model with *δ*_*N*_ (*T*) applied to (9). From Figure 8(c), we observe the emergence of a chronic wound as we expect, displaying consistent qualitative features as in Figure 7(c) when considering increasing *δ*_*N*_. For *β*_1_ *< δ*_*N*_ *< β*_3_, i.e. *δ*_*N*_ ∈ *I*_2_, *I*_3_, however, we observe another partially-healed wound in the travelling wave simulations in Figure 8(b) in contrast to Figure 7(b) and demonstrates the existence of two qualitatively different travelling wave profiles, depending on the choice of initial conditions. We observe from Figure 8(a) that for a chronic wound to heal, *δ*_*N*_ (*t*) must be decreased to when *δ*_*N*_ *< β*_1_, i.e *δ*_*N*_ ∈ *I*_1_, demonstrating the subcritical nature of the bifurcation point *β*_1_, and the multistable nature of different travelling wave profiles for *β*_1_ *< δ*_*N*_ *< β*_3_. The multistability of the travelling wave profiles occurs as there is history dependence upon increasing and decreasing the apoptotic rate, i.e. a hysteresis loop affects the invading states of the travelling waves profiles. We conclude that chronic wounds may persist if apoptotic rates reach an unregulated state, but these chronic wounds may heal to completion if *δ*_*N*_ were able to be regulated from a deregulated state, for example by controlling blood sugar levels.

**Figure 8:**
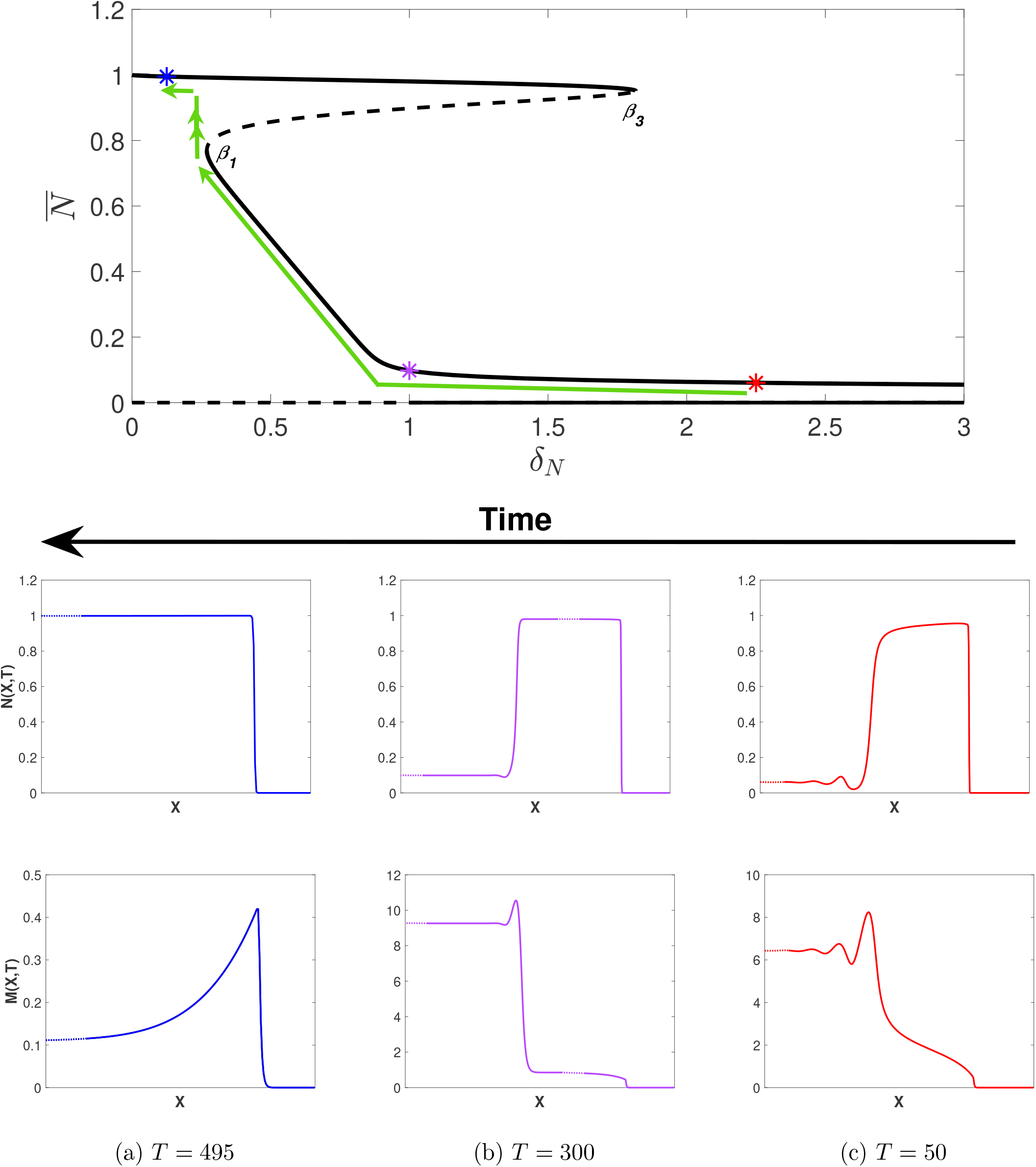
Simulations of the wound healing model with (24) and *ϵ* → −*ϵ* applied to (9), i.e. *δ*_*N*_ decreasing in time with 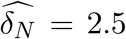 and *ϵ* = 0.005. Other parameter values are as given in Table 2. Note that the *X*-axis has been trimmed to highlight the qualitative features of each travelling wave (dotted lines). The bifurcation diagram is as given in Figure 5(a) for *δ*_*N*_ ∈ [0, 3] and coloured asterisks represent the *δ*_*N*_ (*T*) values at the time points given in (a), (b) and (c). Note that all travelling waves shown connect to the zero state.

## 6 Elevated MMP Production

As described in Section 1, chronic wounds are characterised by elevated levels of MMPs which prevent its healing. MMP production is a factor that may potentially be targeted with treatments for chronic wounds. In a similar fashion to the varying of *δ*_*N*_, we now examine the qualitative changes to travelling wave profiles when we vary the production rate of MMPs, *α*.

We present bifurcation diagrams for 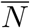 and 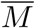 against *α* in Figure 9; the stability of the branches is determined by linear stability analysis detailed in Appendix B. Inspection of the bifurcation diagrams show that we have two saddle-node bifurcation points (*μ*_1_ and *μ*_2_) separating three parameter regimes. We note that at *μ*_2_ (at which *α* is large), the branch of healed states of 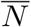 is destroyed and hence, similar to in Section 4.3, we expect the emergence of a chronic wound.

**Figure 9:**
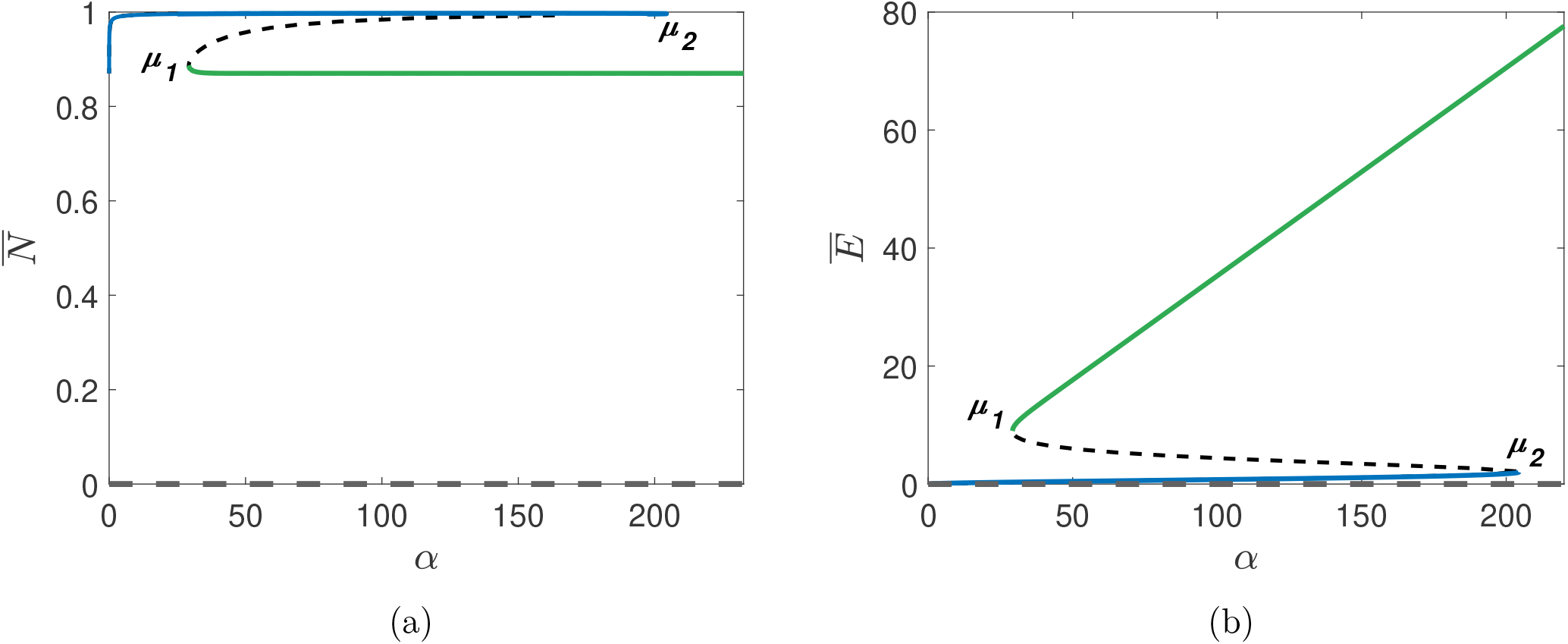
Bifurcation diagrams of *α* against (a) 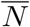, and (b) 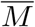. Other parameter values are as given in Table 2. The stable branches are colour-coded such that the green branch of steady states of 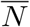 in (a) correspond to the green branch of steady states of 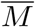 in (b); equivalent for blue branches.

Similar to Section 5, we now investigate the dynamics of the wound healing model for both increasing and decreasing *α*. Firstly, we allow *α* in (10) to slowly increase in time:

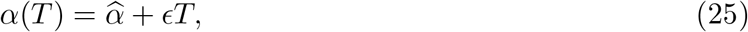

where 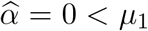. As we see in Figure 10, the travelling wave for *N*(*X, T*) transitions from a completely healed wound to a partially healed wound with elevated MMP concentrations. These dynamics occur as *α* exceeds *μ*_*_ ≈ 103.2 where, despite the invading state of the travelling wave attaining a healed value, the wave visits the unhealed state, resulting in a partially healed wound. Increasing *α* further results in the the invading state transitioning to an unhealed value, and thus the emergence of a chronic wound at *μ*_2_. We deduce that the threshold MMP production rate at which wounds cease to heal to completion is *μ*_*_, rather than the saddle-node bifurcation point *μ*_2_ as we may have originally expected.

**Figure 10:**
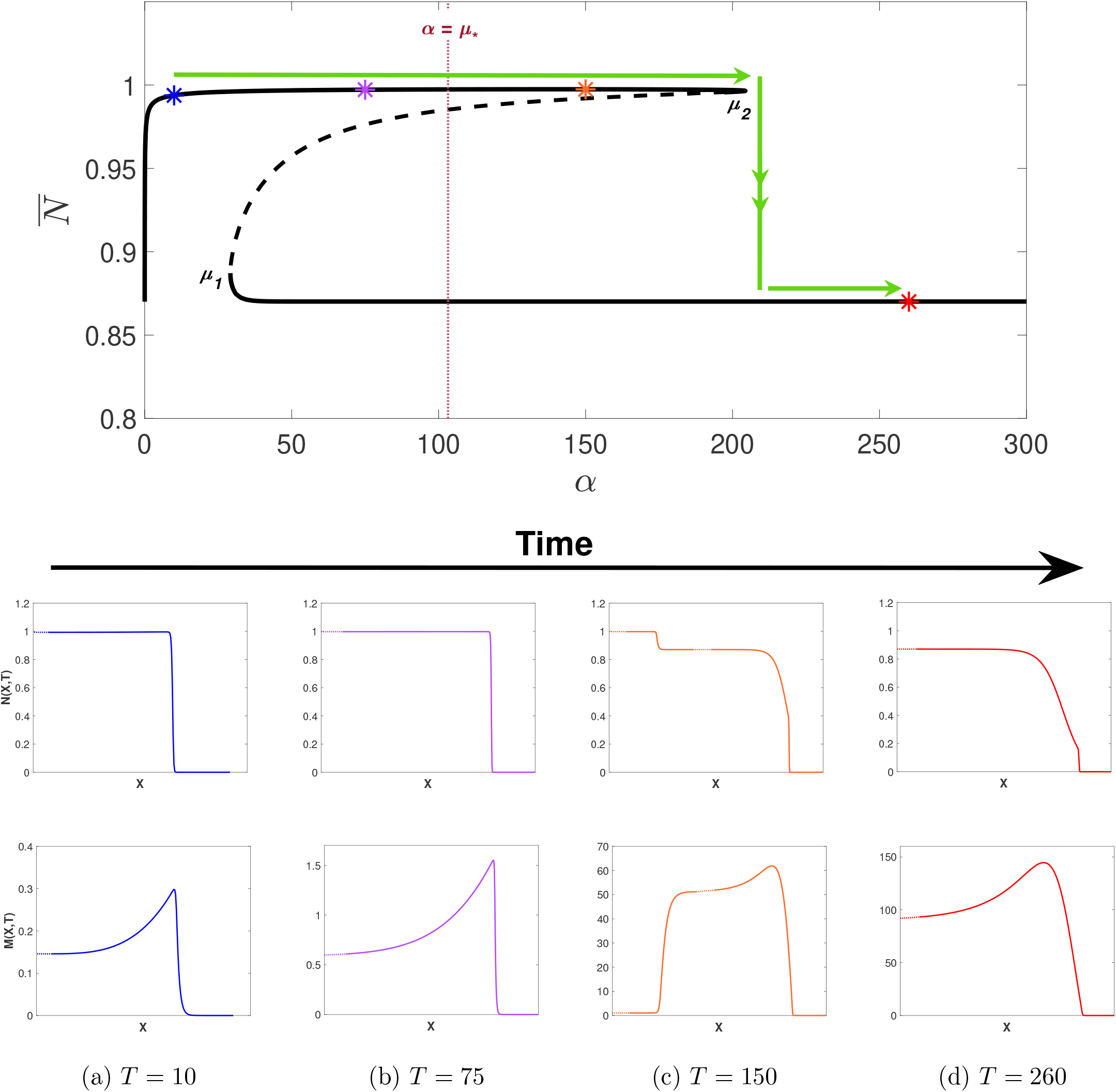
Simulations of the wound healing model with (25) applied to (9), i.e. *α* increasing in time with 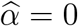 and *ϵ* = 1. Other parameter values are as given in Table 2. Note that the *X*-axis has been trimmed to highlight the qualitative features of each travelling wave (dotted lines). The bifurcation diagram is as given in Figure 5(a) for *α* ∈ [0, 300] and 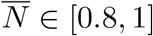 and coloured asterisks represent the *α*(*t*)) values at the time points given in (a), (b), (c) and (d). The maroon dotted line represents the value *α* = *μ*_*_. Note that all travelling waves shown connect to the zero state.

We now consider decreasing *α* to investigate whether not a chronic wound may be reversed if MMP production levels were able to be regulated. In Figure 11, we take *α* = 300 *> μ*_2_ and allow *α* in (10) to slowly decrease in time via (25) by changing *ϵ* with −*ϵ*. From Figure 11(d), we observe the emergence of a chronic wound as we expect, showing consistent qualitative features as in Figure 10(d) when considering increasing *α*. For *μ*_1_ *< α < μ*_2_ however, we observe wounds that do not heal to completion in contrast to the case of increasing *α*, again demonstrating the existence of multistable travelling wave profiles depending on the choice of initial conditions. We also observe a change in qualitative features at *α* = *μ*_*_. For *α < μ*_*_, the travelling wave visits the healed state resulting in a partially healed wound. From Figure 11(d), we deduce that *α*(*T*) must be reduced such that *α < μ*_1_ for a chronic wound to heal.

**Figure 11:**
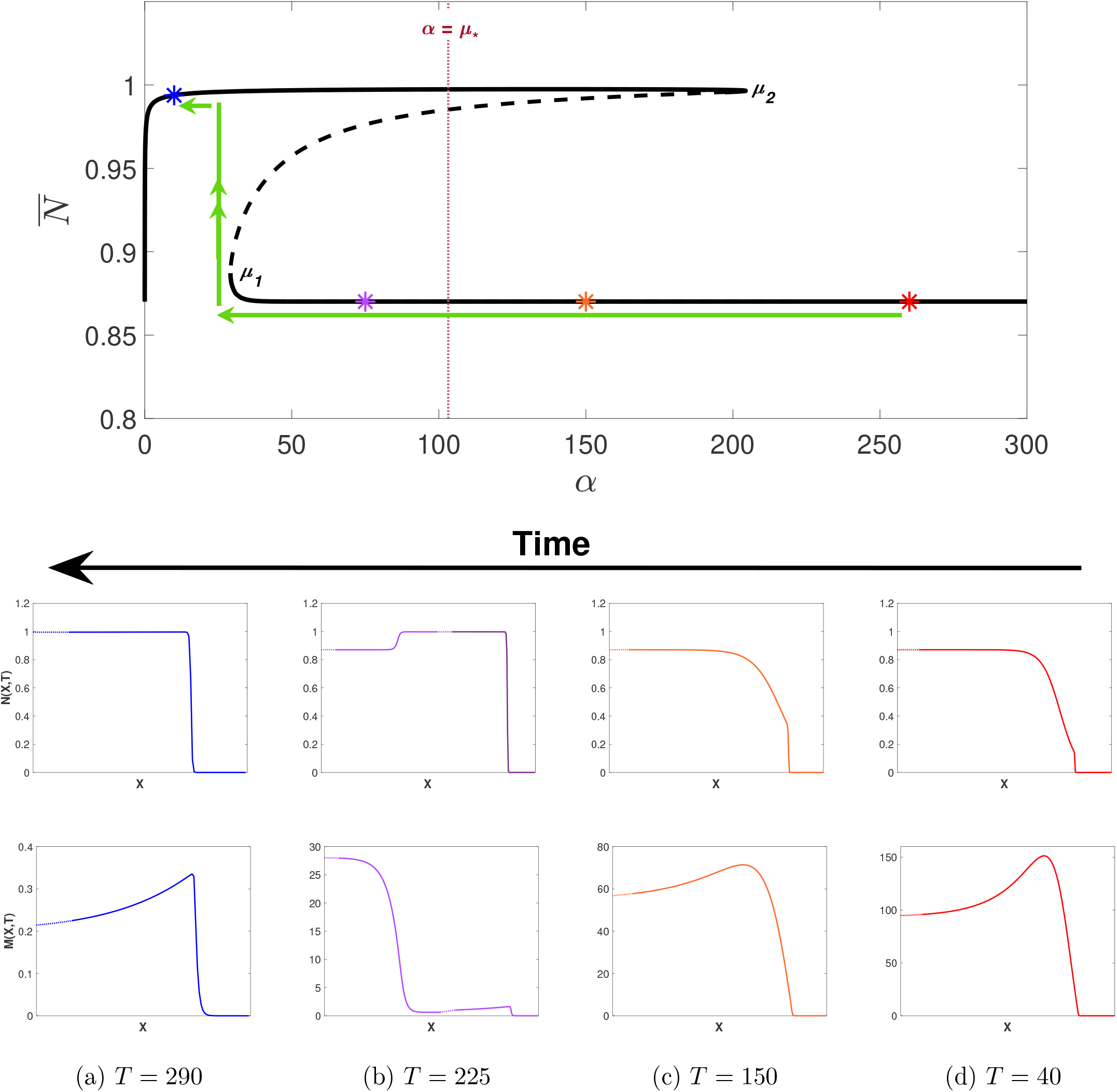
Simulations of the wound healing model with (25) and *ϵ* → −*ϵ* applied to (9), i.e. *α* increasing in time with 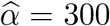 and *ϵ* = 1. Other parameter values are as given in Table 2. Note that the *X*-axis has been trimmed to highlight the qualitative features of each travelling wave (dotted lines). The bifurcation diagram is as given in Figure 5(a) for *α* ∈ [0, 300] and 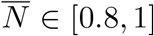 and coloured asterisks represent the *α*(*t*)) values at the time points given in (a), (b), (c) and (d). The maroon dotted line represents the value *α* = *μ*_*_. Note that all travelling waves shown connect to the zero state.

Similar to in Section 5, there is a hysteresis loop which results in history dependence of the travelling wave profiles on increasing and decreasing *α* which results in multistable travelling wave profiles for *μ*_1_ *< α < μ*_2_. Unlike in Section 5 however, we observe an inconsistency in qualitative features of travelling waves for *μ*_1_ *< α < μ*_2_ due to the existence of the threshold value *μ*_*_. We conclude that that wounds will only partially heal beyond *α* = *μ*_*_. Furthermore, a chronic wound may be reversed to a healed state if MMP production rate is controlled from a deregulated state back to a regulated state.

## 7 Conclusions and Discussion

In this work, we develop a two-variable reaction-diffusion model, describing the interaction of matrix metalloproteinases (MMPs) with dermal cells in the wound healing process. In particular, we focus attention on the emergence of chronic wounds, since biological literature suggests that elevated levels of MMPs play a key role in their emergence. Our mathematical model gives rise to travelling wave solutions and in particular is able to emulate key qualitative features of the wound healing process reported in biological literature. One such property is that under parameter regimes representing healthy biological functioning, we observe acute wounds healing to completion with an increased expression in MMP concentration at the edge of the healing wound.

To contrast acute wounds with chronic wounds, we consider the effect of varying the apoptotic rate of dermal cells. Small apoptotic rates correspond to healthy biological functioning and thus we observe the complete healing of a wound. Provided initial concentrations of MMPs are kept low, an increase in apoptotic rate also results in the healing of a wound, which suggests that in order to avoid risk of the emergence of a chronic wound, baseline levels of MMPs must be kept minimal. Elevated apoptotic rate beyond a threshold value leads to the emergence of a chronic wound, with elevated levels of MMPs invading the wound which prevents its healing. Moreover, we observe mulitstable travelling waves depending on initial conditions due to the existence of a hysteresis loop in the bifurcation diagrams for apoptotic rate. Therefore, in order to reverse a chronic wound to a state of complete healing, the apoptotic rate must be decreased below a threshold value. This give insights into the regulation of apoptotic rate in the healing of chronic wounds which may be achieved by, for example, regulating blood sugar levels in diabetes patients.

We also consider the effect of varying MMP production rate on the healing of a wound. Similar to the analysis for apoptotic rate, we obtain threshold values at which chronic wounds persist and may be reversed. However, contrasting from the variation of the apoptotic rate, we observe a change in the the qualitative features of the travelling wave solutions at a different threshold value *μ*_*_. At this new threshold, occurring in the middle of the multistable regime, a wound transitions from healing completely to only being partially healed, before reaching a regime where chronic wounds persist.

Due to the observation of multistable travelling wave solutions with varying qualitative features, a natural extension to this work includes conducting a stability analysis of these travelling waves. Furthermore, despite the wound healing model successfully being able to emulate qualitative features of both acute and chronic wounds, it has greatly simplified the process of wound healing by coupling the action of fibroblasts and ECM, as well as assumed that MMPs directly contribute to healing of cells. In reality, MMPs indirectly contribute to the healing by allowing the migration of other key elements involved in the wound healing process, such as fibroblasts, keratinocytes and endothelial cells [3]. Future research should therefore take these factors into consideration. Furthermore, the role of tissue inhibitors of MMPs (TIMPs) in the wound healing process should be considered as these have an impact on the regulation of MMP concentration. This consideration would ultimately give further insight into the behaviour of MMPs during the wound healing process, as well as potentially direct biological therapies of chronic wounds, i.e. by directing the optimum physical and chemical composition of hydrogel therapies.

## Appendix A: Alternative *f*(*m*)

In Section 2, we consider the function form of *f*(*m*) in (1) which describes the healing contribution of dermal cells by MMPs. Since large concentrations of MMPs results in the destruction of the ECM, we may expect *f*(*m*) to become negative for large *m*. One such function that captures this behaviour is the following:

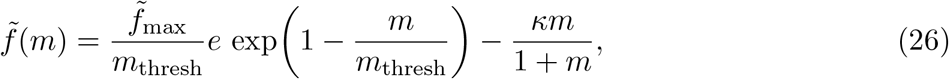

for some values of 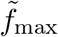 and *κ*. A schematic of 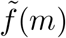 is shown in Figure 12. In non-dimensionalising the PDE system (1)–(2) subject to 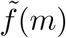 given in (26), using the scaling given in (8), we obtain the simulations given in Figure 13. Initial and boundary conditions are chosen as in (12)–(14) and we take 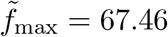 and *κ* = 20; all other parameter values are as given in Table 1. In Figure 13, we observe largely similar qualitative features as those seen in Figure 3: the invasion of a wound by a healed front of cells with an increased expression of MMP concentration at the wound edge.

**Figure 12:**
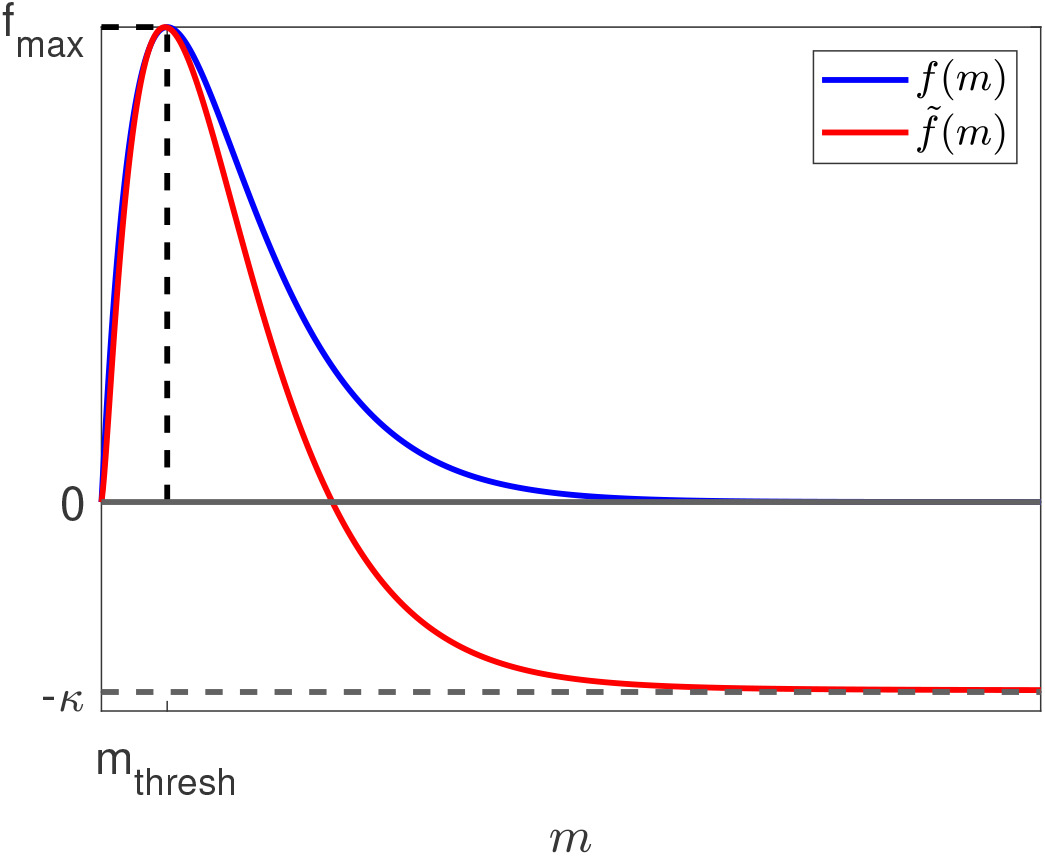
Schematic function forms of the healing contribution by MMPs functions ***f*** (*m*) shown in (2a) and 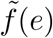 shown in (26)

**Figure 13:**
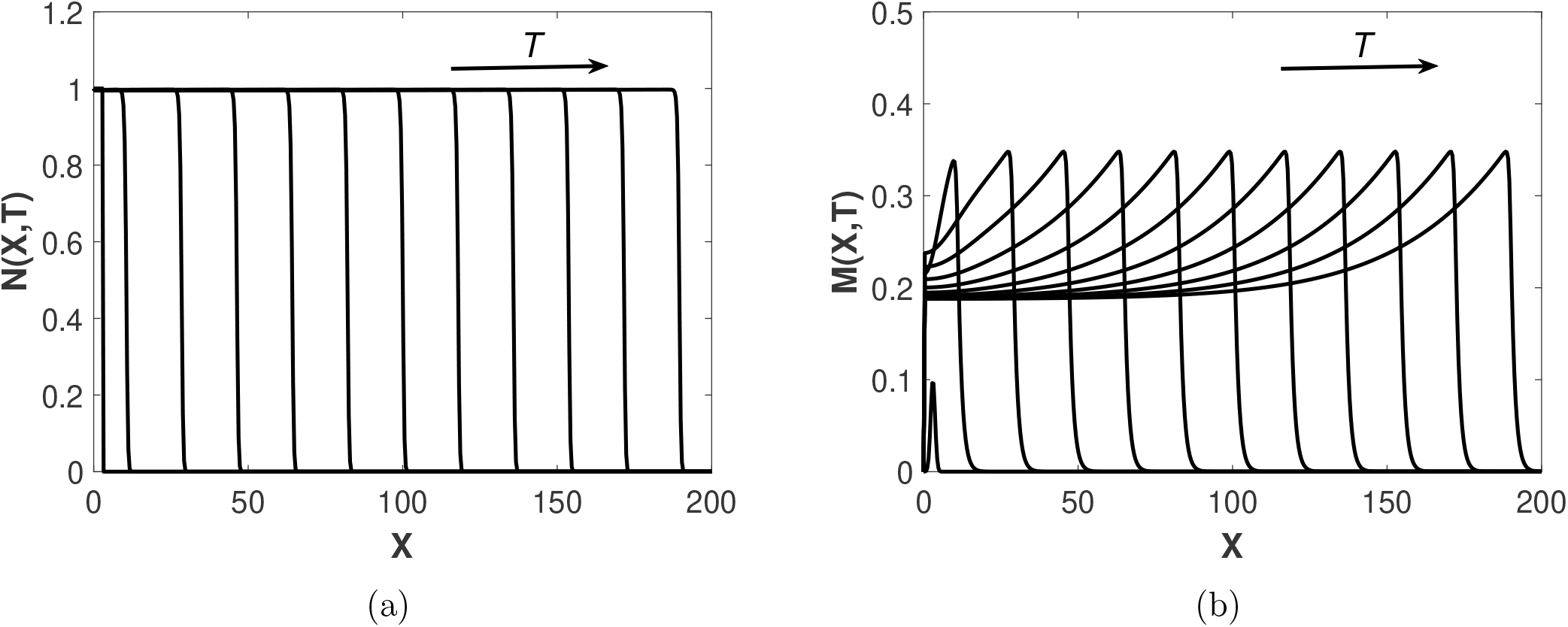
Simulation of the system (1)–(7), non-dimensionalised according to (8) with *f*(*m*) as in (26) at regular time intervals with 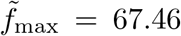, *κ* = 20 and other parameter values as in Table 2. (a) Evolution of cell density *N*(*X, T*) across the spatial domain, (b) Evolution of MMP concentration *M*(*X, T*) across the spatial domain.

## Appendix B: Steady State Stability Analysis

In this section, we consider the stability of spatially-uniform steady states that solve (18)–(19). We note that the Jacobian of this system is given by:

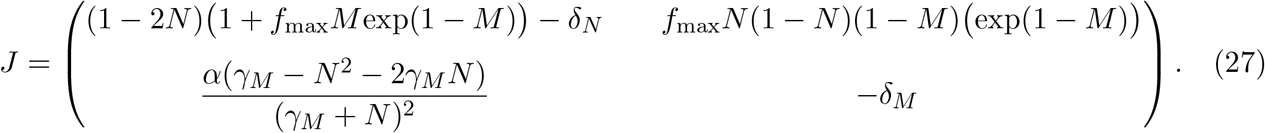

At the steady state 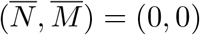, the Jacobian becomes

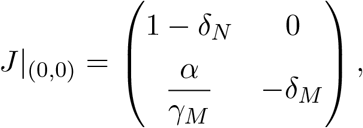

which provides the following eigenvalues:

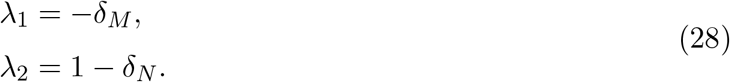

Since *δ*_*M*_, *δ*_*N*_ *>* 0, the zero state is an unstable saddle for *δ*_*N*_ *<* 1 and is a stable node for *δ*_*N*_ *>* 1. For all other steady states, we use MATLAB’s fsolve function to determine 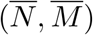 which are then substituted into (27) and whose eigenvalues are computed using MATLAB’s eig function.

## Appendix C: Bifurcation Points *δ*_*N*_

From the bifurcation diagrams in Figure 5, we observe that there are four bifurcation points: one transcritical and 3 saddle nodes. The value of *δ*_*N*_ at which the transcritical bifurcation point occurs may be deduced from the argument in Appendix B. From (28), we conclude that the zero state is an unstable saddle for *δ*_*N*_ *<* 1 and is a stable node for *δ*_*N*_ *>* 1. This change in stability confirms that the transcritical bifurcation point occurs at 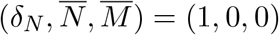.

The values of 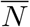 at which the saddle node bifurcation points occur may be calculated by computing the derivative of (17) with respect to 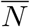 and setting this equal to zero, i.e.

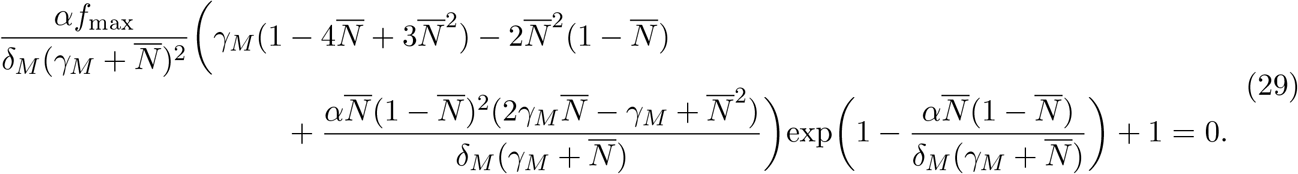

We use MATLAB’s fzero function to find the 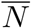 values of the saddle node bifurcation points, and then substitute these into (15) and (17) to find the corresponding 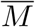 and *δ*_*N*_ values respectively. For the parameter choices as in Table 2, the bifurcation points are 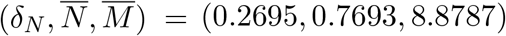, (1.8164, 0.9508, 2.0465) and (50.5904, 0.007998, 0.9916). I.e. the values of *β*_1_, *β*_2_, *β*_3_ and *β*_4_ as discussed in Section 4, subject to other parameter values as in Table 2, are *β*_1_ = 0.2965, *β*_2_ = 1, *β*_3_ = 1.9164 and *β*_4_ = 50.5904.

## References

[1] H.A. Wallace, B.M. Basehore, and P.M. Zito. Wound Healing Phases. StatPearls Publishing, 2021. doi: https://www.ncbi.nlm.nih.gov/books/NBK470443/.

[2] P Martin. Wound healing - aiming for perfect skin regeneration. Science, 276(5309):75–81, 1997.

[3] T.T Nyugen, S Mobashery, and M Chang. Roles of Matrix Metalloproteinases in Cutaneous Wound Healing. Intech Open, 2016.

[4] F Fan, S Saha, and D Hanjaya-Putra. Biomimetic hydrogels to promote wound healing. Front. Bioeng. Biotechnol., 9(718377):1–24, 2021.

[5] Department of Health and Social Care. Advanced wound care: develop new treatments in the uk. URL https://www.gov.uk/government/publications/advanced-wound-care-develop-treatments-in-the-uk/advanced-wound-care-develop-new-treatments-in-the-uk.

[6] R.G Frykberg and J Banks. Challenges in the treatment of chronic wounds. Adv Wound Care New Rochelle, 4(9):560–582, 2015.

[7] A.J Flegg, S.N Menon, P.K Maini, and D.L.S McElwain. On the mathematical modelling of wound healing angiogenesis in skin as a reaction-transport process. Front. Physiol., 6(262): 1–17, 2015.

[8] C Frantz, K.M Stewart, and V.M Weaver. The extracellular matrix at a glance. J Cell Sci., 123(24):4195–4200, 2010.

[9] S Guo and L.A DiPietro. Factors affecting wound healing. J Dent Res, 89(3):219–229, 2010.

[10] A Krejner, M Litwiniuk, and T Grzela. Matrix metalloproteinases in the wound microenvironment: therapeutic perspectives. Chronic Wound Care Management and Research, 3:29–39, 2016.

[11] M Li, A Moeen Rezakhanlou, C Chavez-Munoz, A Lai, and A Ghahary. Keratinocyte-releasable factors increased the expression of mmp1 and mmp3 in co-cultured fibroblasts under both 2d and 3d culture conditions. Mol Cell Biochem, 332(1-2):1–8, 2009.

[12] F Sabino and U.A.D Keller. Matrix metalloproteinases in impaired wound healing. Metalloproteinases In Medicine, 2:1–8, 2015.

[13] N.K Rai, K Tripathi, D Sharma, and V.K Shukla. Apoptosis: A basic physiologic process in wound healing. The International Journal of Lower Extremity Wounds, 2(3):138–144, 2005.

[14] A.K Arya, R Tripathi, S Kumar, and K Tripathi. Recent advances on the association of apoptosis in chronic non healing diabetic wound. World J Diabetes, 5(6):756–762, 2014.

[15] T Sun, S Adra, R Smallwood, M Holcombe, and S MacNeil. Exploring hypotheses of the actions of tgf-β1 in epidermal wound healing using a 3d computational multiscale model of the human epidermis. PLoS One, 4(12), 2009.

[16] Q Mi, B Rivi`ere, G Clermont, D.L Steed, and Y Vodovotz. Agent-based model of inflammation and wound healing: insights into diabetic foot ulcer pathology and the role of transforming growth factor-β1. Wound Repair Regen., 15(5):672–682, 2007.

[17] D.C Walker, G Hill, Wood S.M, R.H Smallwood, and J Southgate. Agent-based computational modeling of wounded epithelial cell monolayers. IEEE Trans Nanobioscience, 3(3):153–163, 2004.

[18] C Ziraldo, Q Mi, G An, and Y Vodovotz. Computational modeling of inflammation and wound healing. Adv Wound Care (New Rochelle), 2(9):527–537, 2013.

[19] G An, Q Mi, J Dutta-Moscatto, and Y Vodovotz. Agent-based models in translational systems biology. WIREs Mechanisms of Disease, 1(2):159–171, 2009.

[20] S.C Bankes. Agent-based modeling: A revolution? PNAS, 99(3):7199 –7200, 2002.

[21] J.A Sherratt and J.D Murray. Models of epidermal wound healing. Proc. Biol. Sci., 241(1300): 29–36, 1990.

[22] J.A Sherratt and J.D Murray. Epidermal wound healing: the clinical implications of a simple mathematical model. Cell Transplant, 1(5):365–371, 1992.

[23] P.D Dale, J.A Sherratt, and P.K Maini. The speed of cornea1 epithelial wound healing. Appl. Math. Lett., 7(2):11–14, 1994.

[24] H Sheardown and Y.L Cheng. Mechanisms of corneal epithelial wound healing. Chemical Engineering Science, 51(19):4517–4529, 1996.

[25] E.A Gaffney, P.K Maini, C.D McCaig, M Zhao, and J.V Forrester. Modelling corneal epithelial wound closure in the presence of physiological electric fields via a moving boundary formalism. IMA Journal of Mathematics Applied in Medicine and Biology, 16(4):369–393, 1999.

[26] N.X. Landen, D. Li, and M. Stahle. Transition from inflammation to proliferation: a critical step during wound healing. Cell Mol Life Sci, 73(20):3861–3885, 2016.

[27] M.A.J Chaplain and H.M Byrne. Mathematical modelling of wound healing and tumour growth: two sides of the same coin. wounds. Wounds: a Compendium of Clinical Research and Practice, 8(2):42–48, 1996.

[28] G.J Pettet, M.A.J Chaplain, D.S.L McElwain, and H.M Byrne. On the role of angiogenesis in wound healing. Proc. R. Soc. Lond. B Biol. Sci., 263:1487–1493, 1996a.

[29] G.J Pettet, H.M Byrne, D.S.L McElwain, and J Norbury. A model of wound-healing angiogenesis in soft tissue. Math. Biosci., 136:35–63, 1996b.

[30] R.C Schugart, A Friedman, R Zhao, and C.K Sen. Wound angiogenesis as a function of tissue oxygen tension: a mathematical model. Proc. Natl. Acad. Sci. U.S.A, 105(2628–2633), 2008.

[31] P.D Dale, J.A Sherrat, and P.K Maini. Role of fibroblast migration in collagen fiber formation during fetal and adult dermal wound healing. Bull. Math. Biol., 59(6):1077–1100, 1997.

[32] Wearing H.J and J.A Sherratt. Keratinocyte growth factor signalling: a mathematical model of derma-epidermal interaction in epidermal wound healing. Mathematical Biosciences, 165 (1):41–62, 2000.

[33] S.N Menon, J.A Flegg, S.W McCue, R.C Schugart, R.A Dawson, and D.S.L McElwain. Modelling the interaction of keratinocytes and fibroblasts during normal and abnormal wound healing processes. Proc. Biol. Sci., 279(1741):3329–3338, 2012.

[34] S.N Menon and J.A Flegg. Mathematical modeling can advance wound healing research. Adv Wound Care (New Rochelle), 10(6):328–244, 2021.

[35] J.C Dallon, J.A Sherratt, and P.K Maini. Mathematical modelling of extracellular matrix dynamics using discrete cells: fiber orientation and tissue regeneration. J Theor Biol, 1999(4): 449–471, 1999.

[36] J.C Dallon, J.A Sherratt, P.K Maini, and M Ferguson. Biological implications of a discrete mathematical model for collagen deposition and alignment in dermal wound repair. IMA J Math Appl Med Biol., 17(4):379–393, 2000.

[37] J.C Dallon, J.A Sherratt, and P.K Maini. Modeling the effects of transforming growth factor-beta on extracellular matrix alignment in dermal wound repair. Wound Repair Regen., 9(4): 278–286, 2001.

[38] S McDougall, J.C Dallon, J.A Sherratt, and P.K Maini. Fibroblast migration and collagen deposition during dermal wound healing: mathematical modelling and clinical implications. Philos Trans A Math Phys Eng Sci, 364(1843):1385–1405, 2006.

[39] B.D Cumming, D.L.S McElwain, and Z Upton. A mathematical model of wound healing and subsequent scarring. J R Soc Interface, 7(42):19–34, 2009.

[40] P.K Maini, D.L.S McElwain, and D Leavesley. Travelling waves in a wound healing assay. Appl. Math. Lett., 17(5):575–580, 2004.

[41] P.K Maini, D.L.S McElwain, and D Leavesley. Travelling wave model to interpret a wound-healing cell migration assay for human peritoneal mesothelial cells. Tissue Engineering, 10 (3/4):475–482, 2004.

[42] L Michaelis and M.L Menten. Die kinetik der invertinwirkung. Biochem. Z, 49:333–369, 1913.

[43] I.E Collier, W Legant, B Marmer, O Lubman, S Saffarian, T Wakatsuki, E Elson, and G.I Goldberg. Diffusion of mmps on the surface of collagen fibrils: The mobile cell surface – collagen substratum interface. PLoS One, 6(9):1–14, 2011.

[44] Y Jin, H Han, J Berger, Q Dai, and M.L Lindsey. Combining experimental and mathematical modeling to reveal mechanisms of macrophage-dependent left ventricular remodeling. BMC Syst Biol, 5(1):60–74, 2011.

[45] N.E Deakin and M.A.J Chaplain. Mathematical modeling of cancer invasion: The role of membrane-bound matrix metalloproteinases. Front Oncol, 3(70):1–9, 2013.

[46] M Muller, C Trocme, B Lardy, F Morel, S Halimi, and P.Y Benhamou. Matrix metalloproteinases and diabetic foot ulcers: the ratio of mmp-1 to timp-1 is a predictor of wound healing. Diabet Med, 25(4):419–426.

